# STAG-CNS: An Order-Aware Conserved Non-coding Sequences Discovery Tool For Arbitrary Numbers of Species

**DOI:** 10.1101/120428

**Authors:** Xianjun Lai, Sairam Behera, Zhikai Liang, Yanli Lu, Jitender S Deogun, James C. Schnable

## Abstract

One method for identifying noncoding regulatory regions of a genome is to quantify rates of divergence between related species, as functional sequence will generally diverge more slowly. Most approaches to identifying these conserved noncoding sequences (CNS) based on alignment have had relatively large minimum sequence lengths (⩾15 base pair) compared to the average length of known transcription factor binding sites. To circumvent this constraint, STAG-CNS integrates data from the promoters of conserved orthologous genes in three or more species simultaneously. Using data from up to six grass species made it possible to identify conserved sequences as short at 9 base pairs with FDP ⩽ 0.05. These CNS exhibit greater overlap with open chromatin regions identified using DNase I hypersensitivity, and are enriched in the promoters of genes involved in transcriptional regulation. STAG-CNS was further employed to characterize loss of conserved noncoding sequences associated with retained duplicate genes from the ancient maize polyploidy. Genes with fewer retained CNS show lower overall expression, although this bias is more apparent in samples of complex organ systems containing many cell types, suggesting CNS loss may correspond to a reduced number of expression contexts rather than lower expression levels across the entire ancestral expression domain.

## Introduction

Mutations accumulate in different parts of the genome at different rates. Protein coding exons tend to have significantly lower rates of nucleotide substitutions than introns or intergenic sequences. The majority of possible substitutions in protein coding sequence usually change the amino acid sequences of the resulting protein. Many such changes in amino acid sequence will have negative effects on the protein. Therefore, many mutations which occur in protein coding sequence are purged from the genome by purifying selection. Protein coding sequences can be said to be functionally constrained, resulting nucleotide substations accumulating more slowly. In both animals and plants, there are islands of non-coding sequence which also exhibit low rates of nucleotide substitution, indicating that these regions are also subject to functional constraint (Hardison et al., 1997; Levy et al., 2001; Kaplinsky et al., 2002; Guo and Moose, 2003). There regions are referred to as conserved non-coding sequences (CNS) (Hardison et al., 1997), or sometimes in the animal literature as conserved non-coding elements (CNE) (Shin et al., 2005). In both animals and plants, CNS have been shown to confer extremely specific spatiotemporal patterns of transcriptional regulation (Shin et al., 2005; Visel et al., 2008; Raatz et al., 2011).

Identification of CNS in animals and plants present very different challenges. In animals, many CNS are large (⩾ 100 bp) (Stephen et al., 2008), while different analyses in plants have primarily identified smaller 15-50 bp CNS (Thomas et al., 2007; Baxter et al., 2012; Turco et al., 2013; Haudry et al., 2013). Various algorithmic approaches are currently employed to identify CNS in plants including both manual (Thomas et al., 2007) and automated (Turco et al., 2013) curation of BLASTN results, global alignments of sliding windows of promoter regions (Baxter et al., 2012), the use of whole genome alignment algorithms (Haudry et al., 2013), alignment-independent detection of enriched IUPAC motifs (De Witte et al., 2015), and searches for biologically defined transcription factor binding sites across orthologous genes (Van de Velde et al., 2016). The majority of these approaches are either based on two-at-a-time sequence alignments (Turco et al., 2013; Baxter et al., 2012; Haudry et al., 2013), and/or do not retain information on conserved microsynteny within the promoter (De Witte et al., 2015; Van de Velde et al., 2016).

Here we describe a new method for CNS detection using suffix tree and maximum flow algorithms (See Methods) which identifies sets of sequences conserved in the same order in the promoters of arbitrarily large numbers of orthologous or paralogous genes. This approach is inherently non-pairwise, which allows researchers to adjust the number of genes compared and significance cutoffs for CNS discovery based on the tolerance of their research program to either false positives or false negatives. The analysis below demonstrates that as greater numbers of species are used in the comparison, CNSs with smaller sizes can be identified while retaining equivalent or better false positive discovery rates.

## Results

### A new approach for fast and accurate CNS identification

A comparative genomics approach combining data on conserved sequences and conserved order was employed identify CNSs within the promoters of syntenic orthologous genes drawn from 2-6 grass species. This approach identifies conserved regions i.e. maximal exact match (MEMs) present in all sequences being analyzed, and then searches for and locates the optimal path including the greatest number of conserved sequences without violations of microsynteny using a weighted acyclic graph (Figure 1A). This approach provides flexibility to identify CNS using data from different numbers of species, or multiple paralogous gene copies from a single family and is runtime efficient, only taking several seconds to complete the computation for one group of orthologous or paralogous genes, and few hours to complete the computation for all syntenic orthologous genes present within a group of species. The software tool (STAG-CNS: Suffix Tree Arbitrary Gene number: Conserved Noncoding Sequence) and its source code are now publicly available at https://github.com/srbehera11/stag-cns.

**Figure 1.**
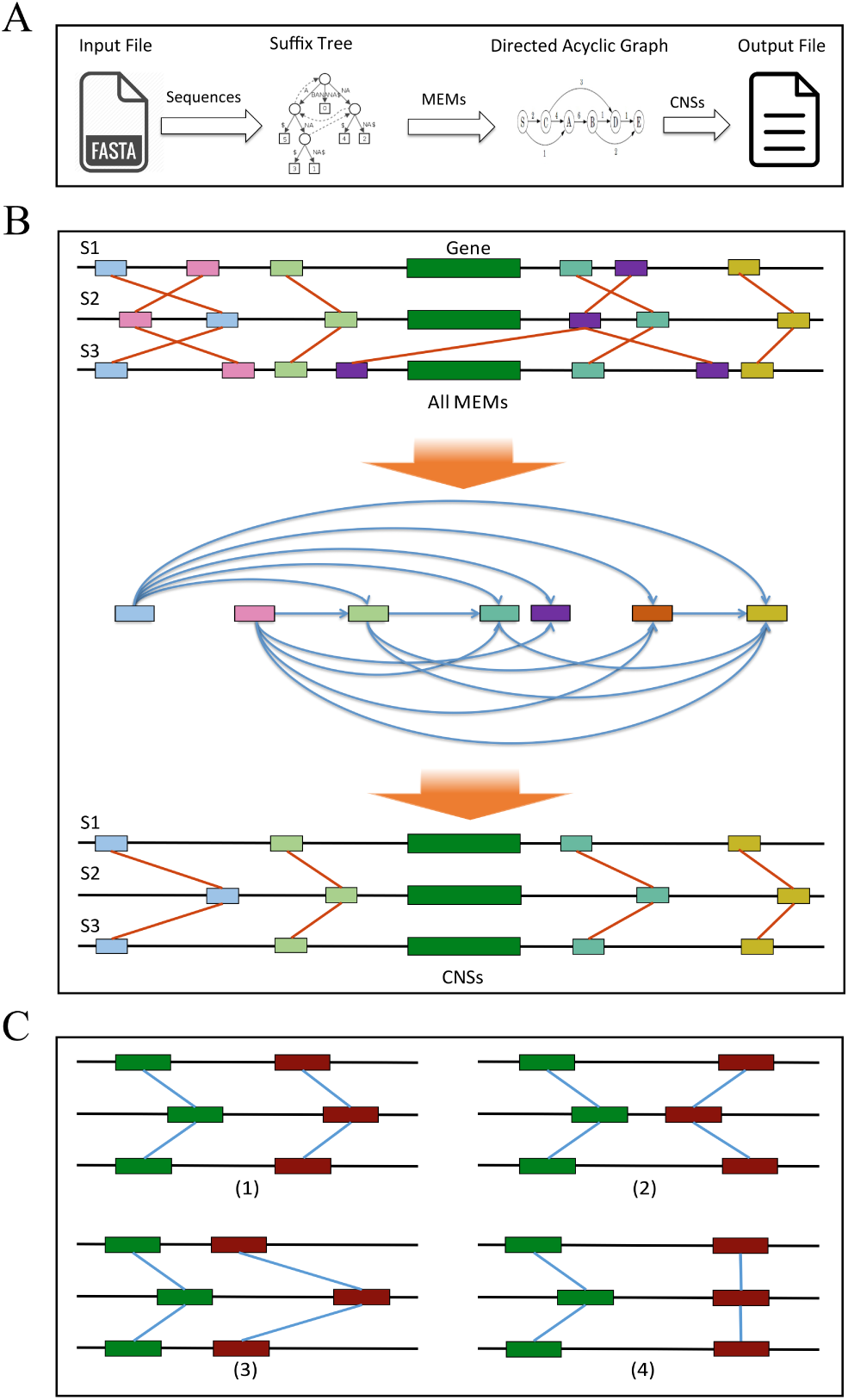
The work-flow for identifying CNS across grass species using STAG-CNS (A) CNS discovery strategy using suffix tree and maximum flow algorithms. (B) CNS of three species and Directed Acyclic Graph using maximal exact matches(MEMs). (C) Breaking ties: Chose the one with highest rank. The ranking of the above scenario is (1)>3)>(4)>(2) (See Methods).

### Estimating the accuracy and sensitivity of STAG-CNS

STAG-CNS has a configurable minimum CNS length, that allows the users balance the trade off between increasing sensitivity to small conserved sequences and controlling false positive discovery rates. The probability of the same short sequence existing in multiple sequences by chance alone is dependent on the number of sequences being compared, the length of each comparator sequence, and the minimum length of the matching sequence. Assuming a random ordering of four nucleotides at equal frequencies this probability can be approximated by the formula: 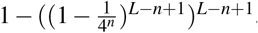, where L represents the length of DNA sequence and *n* means length of small sequence fragment. However, the assumptions given above are violated by most genomes. Among 239 monocot species, GC content was found to vary between 33.6-49.9% (Šmarda et al., 2014). In grasses, individual genes exhibit binomial distributions of gene content (Tatarinova et al., 2010). In addition the frequency of individual short sequences is non-random within a given genome, with certain strings never or rarely occurring (Figure 2A), and others, particularly those found in MITEs and other transposons, occurring at high frequency. Therefore, rather than approximating the false positive discovery rate for STAG-CNS using the formula described above, this parameter was instead estimated using permutation testing, where the number of putative conserved non-coding sequences identified in comparisons of non-orthologous genes was assayed, following the method described by Baxter et al. (Baxter et al., 2012). This method allows the minimum CNS which can be detected at an acceptable false positive rate to be determined empirically. A set of 200 genes (Table S1) conserved at syntenic orthologous located in sorghum, rice, setaria, brachypodium, oropetium, and dichanthelium were randomly selected from a previously published syntenic gene list (Schnable et al., 2016). Conserved non-coding sequences were identified between syntenic orthologous genes in three species – sorghum, rice, and setaria – using minimum CNS lengths between 8 and 22 base pairs. Average false positive discovery rates per gene were estimated using 100 random permutations of the dataset (Figure 2B). As expected, both true positives and false positives declined as higher minimum CNS lengths, while the ratio of true positives to false positives increased. Controlling the false positive discovery rate at 5% of total discovered CNS indicated CNS as short at 12 base pairs can be identified with high confidence using data from three species.

**Figure 2.**
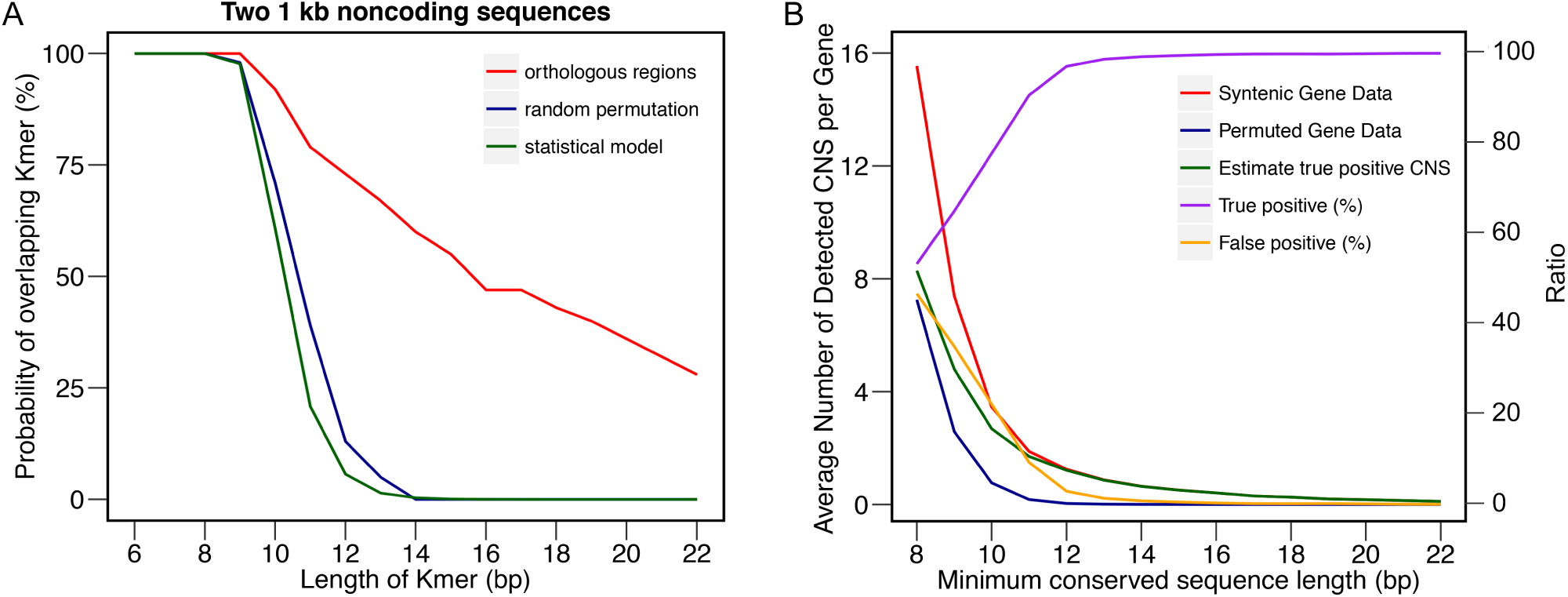
A) The number of overlapping sequences of a given length between two noncoding sequences based on either a statistical model assuming random sequence and equal frequencies of all four nucleotides, random regions extracted from actual grass genomes, or syntenic orthologous non-coding regions extracted from grass genomes. B) Relationship between the minimum length of shared subsequence before it is considered a CNS, number of CNS discovered, and false discovery percentage.

To determine how minimum CNS length responded to variation in the number of species employed in the comparison, CNS were identified between the same set of 200 genes employed above (Table S1) using data from syntenic gene copies in 2, 4, 5, or 6 species, with minimum CNS lengths from 8 to 22 base pairs (Figure 3A). The corresponding false positive discovery rates are displayed in Figure 3B. As expected, increasing the number of species included in the analysis improves the proportion of true positive CNS identified at any given minimum CNS length and with six species it was possible to identify conserved sequences as short as 9 base pairs while maintaining a false discovery rate less than 5%.

**Figure 3.**
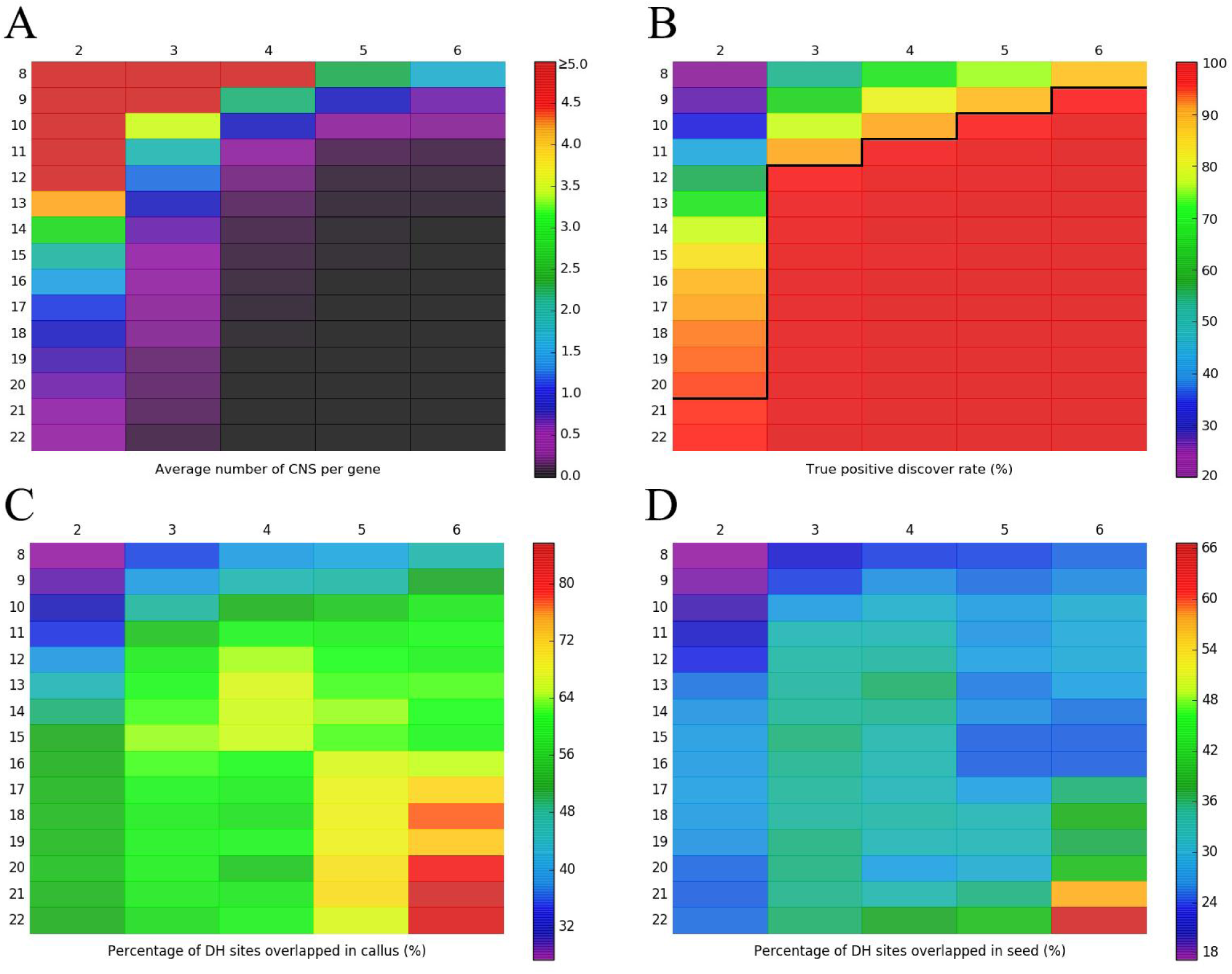
Analysis showing that both increasing the number of species and rising the length of minimum can increase the power of STAG-CNS to detect the CNS. For each sub-figure the x-axis shows the species number analyzed and y-axis shows the minimum length used to identify CNS. (A) The number of CNS identified in different number of species and the minimum length of CNS. (B) True positive discovery rate of CNS across different number of species. The true positive rates below the black line indicated reliable minimum length of CNS (true positive rates ¿=95%). (C) and (D) Overlap rate of CNS and DNase sites in callus and seed.

### Comparison of results from STAG-CNS and CDP

The CNS Discovery Pipeline (CDP) is one of the tools previously used to identify conserved non-coding sequences among different grass species (Turco et al., 2013). Unlike STAG-CNS, the CDP works by performing pairwise comparisons based on BLASTN, and identifies CNS present in three or more species through overlap with a single common reference. The CNSs were identified between 200 genes conserved in sorghum, setaria, and rice using the CDP and STAG-CNS for direct comparisons of the results produced by both methods. A set of genes previously shown to be CNS rich were selected to maximize the number of informative comparisons (Table S3). The CDP was run with the default minimum CNS length of 15 bp for sorghum-rice and sorghum-setaria comparisons. A CNS was considered to be present across all three species if there was at least a 12 bp overlap in sorghum between a sorghum-rice CNS and sorghum-setaria CNS. The number and total length of three-species CNS identified for each syntenic orthologous gene group was compared between the two methods and found to be moderately correlated. Pearson correlation coefficient of CNS number and total length of CNS between two methods was 0.415 (p-value = 1.01e-09) and 0.527 (p-value = 1.332e-15), respectively (Figure S1).

Subsequently, all of the 17,996 orthologous syntenic genes of three species (sorghum, setaria, and rice) (Table S2) were used to identify the CNS using STAG-CNS and CDP (sorghum and rice, sorghum and setaria, and then the pan-grass species). The CNS information from both methods was summarized (Table 1) and the orthologous CNS of the syntenic genes of three species were listed in Table S4. The CDP uses nucleotide-nucleotide BLAST (blastn) as its core aligner, allowing it to identify longer sequencers with multiple mismatches or gaps as CNS, while STAG-CNS currently requires exact matches. The mismatch rates for each pairwise CNS identified by the CDP were 9.3% between sorghum and rice and 11.6% between sorghum and setaria, suggesting one mismatch for each 10 bp sequences on average in CDP method. As a result STAG-CNS identified fewer CNS per gene in this three-way comparison. Across the 17,996 gene syntenic gene triplets employed in this analysis STAG-CNS identified an average of 0.58 CNS per gene while the CDP identified an average of 1.25 CNS per gene. Both the average length and the median length of CNS identified by STAG-CNS were smaller than that by CDP. Overall, the total length of the conserved non-coding sequence of syntenic genes identified by CDP and STAG-CNS were 774.2 kb and 162.3 kb,respectively.

**Table 1.**
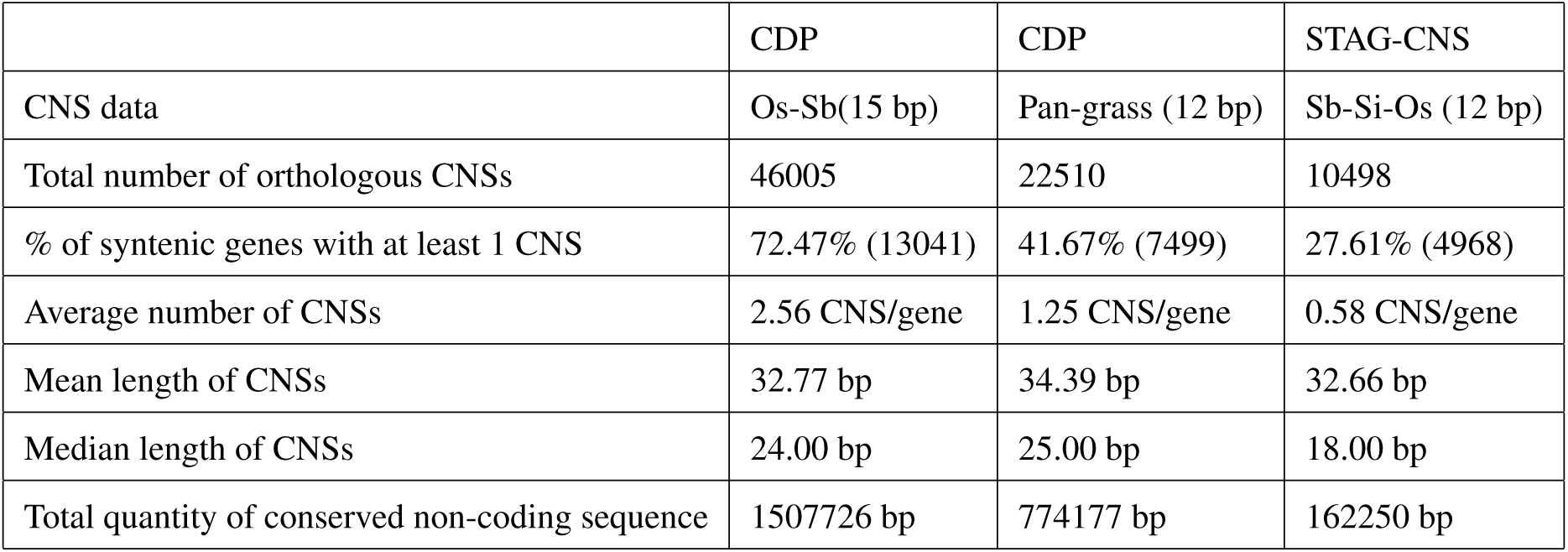
Summary of CNS distributions in 17,996 syntenic orthologous genes.

The CNSs identified by each method were manually proofed for a number of individual syntenic genes. One example (Sobic.008G147500) is shown in (Figure 4). The CDP identified a total of 12 CNSs between sorghum and rice and 8 CNSs between sorghum and setaria. Comparing all three species simultaneously using STAG-CNS identified thirteen short CNSs mostly representing more precise regions within larger CNS identified in pairwise species analysis. Including data from syntenic orthologs in all six grass species employed in this analysis increased the number of CNS identified by STAG-CNS to 18.

**Figure 4.**
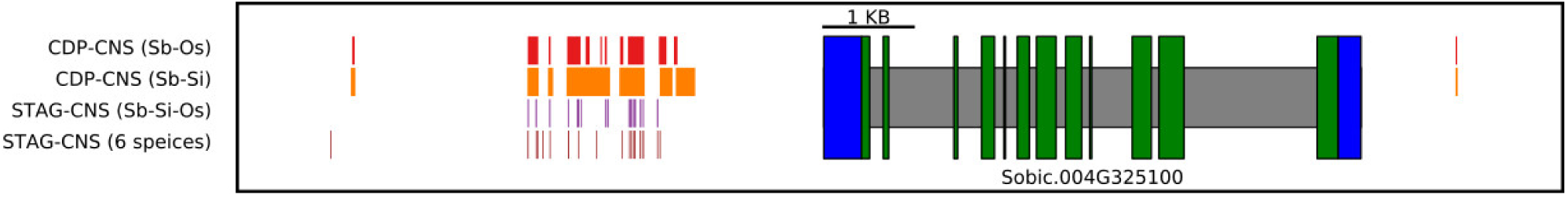
Visual comparison of CNS identified for a given CNS-rich gene using the CDP or STAG-CNS. CNS between Sb-Os and Sb-Si were identified using CDP with default parameters (minimum length of CNS: 15 bp). CNS in Sb-Si-Os was identified using STAG-CNS with a minimum CNS length of 12 bp. CNS in 6 species (Sb-Si-Os-Or-Br-Do) was identified using STAG-CNS with minimum CNS length of 9 bp. The blocks in blue, green, and grey correspond to UTRs, exons, and introns,respectively.

### Association of CNSs with DNase hypersensitive sites

Regulatory sequences are often correlated with regions of open or accessible chromatin (Tsompana and Buck, 2014; Rodgers-Melnick et al., 2016). Open chromatin can be assayed in a whole genome fashion using a range of techniques including FAIRE-seq, MNase-seq, and DNase1 hypersensitivity-seq (Zhang et al., 2012; Rodgers-Melnick et al., 2016). The overlap between CNS identified by STAG-CNS and open chromatin regions was tested using a pre-existing set of DNase hypersensitive sites (DH sites) generated from rice seedling and callus tissue (Zhang et al., 2012). A total of 8,934 CNS identified from the syntenic genes in sorghum, rice, and setaria with the minimum CNS at 12 bp were compared with the DH sites (Figure 5 and Table S5).

**Figure 5.**
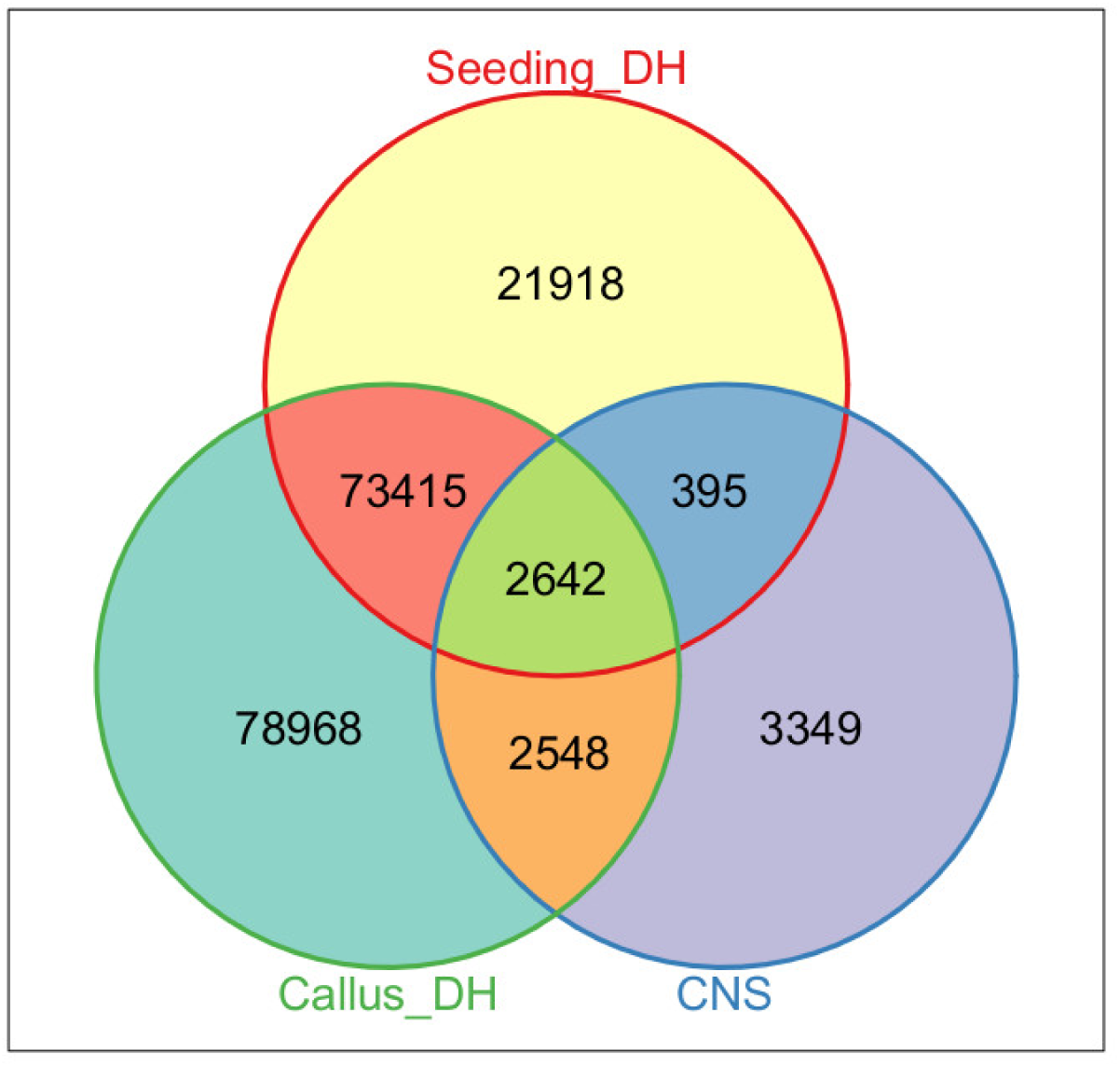
Overlap of rice, sorghum, setaria CNS identified using STAG-CNS with regions of open chromatin identified using DNase1 hypersensitivity in rice seedings and rice callus.

Of these CNS, 34.0% (3,037) and 58.1% (5,190) overlapped with DH sites identified in seeding and callus tissues respectively, significantly more than would be expected from random sequence (binomial test p-value ⩽ 1.0e-06 in both cases). These overlaps are also significantly higher than those previously observed when comparing CNS identified by the CDP between rice and sorghum to the same rice open chromatin datasets (25.7% and 41.6% for seedling and callus tissue respectively) (Zhang et al., 2012).

A set of 1,873 rice genes with conserved syntenic orthologs across all 6 species using in this analysis were used to test how the relative overlap between STAG-CNS identified sequences and DH open chromatin regions responded to variation in the number of species employed to identify CNS and minimum CNS length. The overlap rate between these two orthogonal methods of identifying regulatory sequences increases both when increasing the number of species used in the analysis and with increasing minimum CNS length (Figure 3C-D).

### Functional enrichments among CNS rich genes

Previous studies have shown that genes with regulatory functions tend to be associated with greater numbers of CNS (Freeling and Subramaniam, 2009; Turco et al., 2013) and that genes with more complex regulatory patterns tend to have larger promoters in both animals (Nelson et al., 2004) and plants (Sun et al., 2010). Here, genes were grouped based on the number of CNS for each syntenic gene (0, 1, 2, or ⩾ 3). Gene Ontology (GO) enrichment analysis was performed on each group of syntenic genes independently (Table S6). For the genes present in the group with no CNS, nine GO terms related to ‘metabolic process’, ‘catalytic activity’, ‘single-organism metabolic’, and ‘oxidoreductase activity’ etc. were identified as enriched (bonferroni test p-value ⩽ 0.05, FDR ⩽ 0.05). In the group of genes associated with one CNS, 20 GO terms were significantly enriched (bonferroni test p-value ⩽ 0.05, FDR ⩽ 0.05), among these terms 17 were related to ‘regulation of’ and 2 were related to ‘transcription factor activity’ and one was related to ‘biological regulation’. Similarly, 22 GO terms were significantly enriched in the group of genes with two CNSs (bonferroni test p-value ⩽ 0.05, FDR ⩽ 0.05), including all 20 GO terms identified as enriched among genes with one CNS and 2 others related to ‘DNA binding’ and ‘nucleic acid binding’. Among the genes with at least 3 CNSs, a total of 32 GO terms were significantly enriched (bonferroni test p-value ⩽ 0.05, FDR ⩽ 0.05). Among these GO terms, 18 were related to ‘regulation’ and 10 were related to ‘binding.’ Overall these findings are consistent with previous reports using CNS identified using other methods.

A significant proportion of syntenic genes were not associated with any GO term annotations (6,121 genes). The distribution of these unannotated genes across the four categories of CNS-richness described above was tested using a “dummy” GO term not associated with any genes. Significantly fewer genes with no GO annotations were present in the 0 CNS/gene category than expected, and significantly more genes without any GO terms were present in the 1 CNS/gene category. No significant difference from the null model was observed for these genes in the 2 CNS/gene category and ⩾3 CNS/gene category. However, these were also the two categories with the fewest total genes, reducing statistical power to detect significant enrichment or purification of these genes.

### Biased regulatory sequence loss tracks biased gene loss between maize sub-genomes

STAG-CNS can also be used to identify differences in the loss or retention of conserved non-coding sequences between duplicated genes. Maize experienced a whole genome duplication within the last 5-12 million years, after its divergence from the lineage leading to sorghum (Swigoňová et al., 2004) forming two sub-genomes (maize I and maize II) (Schnable et al., 2011). Following the WGD, duplicate copies of many genes were deleted from one or the other sub-genome, however 4-7,000 duplicate gene pairs are retained on both maize sub-genomes today. Gene copies were more likely to be lost from the maize II sub-genome, while gene copies on the maize I sub-genome tend to be expressed at higher levels than their retained duplicates on the maize II sub-genome (Schnable et al., 2011). Unlike other forms of gene duplication, both copies of a WGD derived gene pair are initially associated with equivalent sets of regulatory sequences, although these sequences can be lost from one or the other gene copy, a process known as fractionation mutagenesis (Freeling et al., 2012).

A set of 6,156 syntenic gene groups including gene copies in setaria, sorghum, maize I and maize II) were using in the following analysis. Using the same sorghum and setaria gene, CNS shared by both retained duplicate maize genes, as well as sequences conserved in both out-group species but retained by only one maize gene copy were identified using STAG-CNS (Figure S2A and Table S7). In approximately half the cases (2,925 gene groups), no CNS were found to be associated with either maize gene copy. In an additional 297 cases, equal numbers of CNS were retained by each gene copy. However, of the remaining 2934 cases, approximately 2/3 (1,925 gene groups) showed greater amounts of conserved non-coding sequence associated the maize I copy of a retained duplicate gene pair, and 1/3 (1,010 gene groups) showed greater amounts of conserved non-coding sequence associated with the maize II gene copy. Bias in CNS loss correlated with bias in expression, with greater bias towards expression of the maize I gene copy in cases where the maize I gene copy retained more CNS, and less or no bias towards greater expression of the maize I gene copy when the maize II gene copy retained more CNS. The strength of this effect was quantified in a range of gene expression datasets generated by multiple research groups Zhang et al. (2017); Chang et al. (2012); Bolduc et al. (2012); Waters et al. (2011); Wang et al. (2009); Chettoor et al. (2014); Davidson et al. (2011); Li et al. (2010). Figure S2B shows the difference in the magnitude of biased expression towards maize I between the group of genes where maize I retained more CNS and the group of genes where maize II retained more CNS. Pollen and anthers showed some of the smallest effects of relative CNS number of bias towards maize1 gene copy expression, while whole leaf/whole seedling/whole root samples showed some the the greatest effects of relative CNS number on bias towards maize1 gene copy expression.

## Discussion

We developed an algorithm and software implementation, STAG-CNS, for the identification of conserved non-coding sequences distributed in the noncoding regions flanking conserved syntenic orthologous genes across multiple species. The proposed approach employs the suffix tree and maximum flow algorithm to identify CNS using both sequence conservation constraints, and constrained by conserved microsynteny constraints in each species. Unlikely previous CNS identification tools based on sequence alignment, STAG-CNS can directly align the promoters from three or more species simultaneously. This flexibility makes it possible to identify shorter, but highly conserved, sequences which could not be identified at acceptable false discovery rates using two-at-a-time approaches to CNS discovery (Figure 3B). Effective use of STAG-CNS requires tuning the minimum CNS length. This will vary based on a number of factors including the number of species included in the analysis, the phylogenetic distance between these species, and the total amount of flanking sequence included on either side of target genes. For these reason, the optimal solution for any gene group of species may be to test the false discovery proportions at different minimum CNS lengths as described above, before choosing a cut off for the final analysis.

Known transcription factor binding sites range in length from 6 bp to 15 bp with average length of 10 bp (Stewart et al., 2012; Tuğrul et al., 2015; Yu et al., 2016), so the advance from identifying CNS ⩾ 15 bp to identifying conserved sequences as short at 9 bp with acceptable (⩽ 0.05) false discovery rates means that it should be possible to identify the most conserved binding sites of a wider range of transcription factors using conserved non-coding sequence analysis. As the number of species with high quality genome sequences in groups such as the grasses, crucifers, and legumes increases, it is anticipated it will be possible to employ STAG-CNS to identify even smaller regions of conserved sequence within gene promoters with acceptable false discovery proportions.

Many conserved non-coding sequences identified with previous methods contained mismatches and indels ( 10% of base pair positions for CDP CNS). It is likely that many of these mismatches do not disrupt the function of the conserved noncoding region. Currently STAG-CNS can only identify exact match sequences. However this are precedents in the literature for adapting suffix tree based algorithms to identify conserved sequences containing up to one mismatch without unmanageable increases in runtime or memory requirements. This can be achieved by extending the longest common substring with k mismatch problem for more than two sequences. Crochemore et al. Crochemore et al. (2006) used suffix tree and reverse suffix tree i.e. suffix tree of sequences in reverse to find the longest repeats with k consecutive mismatches among two strings. A modified version of Crochemore’s algorithm was developed by Flouri et al. Flouri et al. (2015) for finding longest common substring with one mismatch. The above method could be extended for more than two sequences, as employed in STAG-CNS, by using a generalized suffix tree and a reverse generalized suffix tree.

Finally, the ability to analyze promoter sequence surrounding three or more genes at once makes it possible to more accurately study the differential loss of conserved sequences from duplicate genes following whole genome duplicate. Here STAG-CNS was used to demonstrate that the maize sub-genome which lost more genes following whole genome duplicate also has less conserved regulatory sequence associated with retained copies of duplicate gene pairs. Several recent reports have suggested that duplicate maize genes have experienced significant regulatory sub- or neo-functionalization (Hughes et al., 2014; Pophaly and Tellier, 2015; Li et al., 2016). The findings here and elsewhere suggest that genes from the maize II subgenome may have sub-functionalized into more specialized expression domains while maize I gene copies retained broader patterns of expression. This model would be broadly consistent with the findings of (Pophaly and Tellier, 2015) that a large population of maize WGD gene pairs exhibit bidirectional expression divergence when examining expression patterns across specific tissue types, and that these genes are enriched in transcription factors, a class of gene which tends to have large promoters containing many conserved regulatory elements.

## Methods

### STAG-CNS algorithm and implementation

The STAG-CNS uses suffix tree and maximum flow algorithm to discover the CNSs in multiple grass species. Repeated sub-sequences are identified using suffix tree and then the microsyntenic path through the target sequences including the greatest number base pairs of repeated subsequences is identified using maximum flow algorithm. The input for STAG-CNS is a fasta file containing fasta sequences of the area surround each gene to be compared, with additional information such as gene name, start and end position of the gene, chromosome name, direction (‘+’ or ‘−’) and the actual start and end positions in the chromosome. All these information is extracted from the annotation files of corresponding species. If the direction is ’−’ i.e. the gene of interest is on the reverse strand, the reverse complement of sequence is constructed prior to analysis. The generalized suffix tree of all sequences is constructed by using Ukonnen’s linear time on-line algorithm (Ukkonen, 1995). The generalized suffix tree is a tree data structure that stores all suffixes of the sequences in a compressed manner. The suffix of a string is a substring which ends at the last position of the string. The leaf nodes of the generalized suffix tree are labeled with the index of the sequence and the start position of the suffix in that sequence. This data structure has been used in variety of applications in computational biology including: search applications, exact match searches, subsequence composition searches, homology searches, single sequence analysis applications, and multiple sequence analysis applications (Bieganski et al., 1994). The maximal exact match (MEM) is a repeat sub-sequence which can not be extended at both the ends. The generalized suffix tree is used to find the MEMs efficiently by traversing the tree and using some additional information associated with suffix tree data structure such as suffix links. The suffix tree was first studied by GusField (Gusfield, 1997) and it was also used for multiple sequence alignment in MGA tool (Höhl et al., 2002). Our algorithm uses the similar approach to find all MEMs with length greater than or equal to the given minimum length (input parameter) using generalized suffix tree. The implementation of this part of the algorithm (finding MEMs from generalized suffix tree) is similar to splitMEM (Marcus et al., 2014). Instead of obtaining all MEMs, the program only finds the MEMs which are a) present in all sequences, b) not present in genomic regions, c) present in the same side of the gene. A weighted directed acyclic graph is constructed using the MEMs obtained in the previous stage. Each MEM becomes a node in the graph and the weight of the node is the length of that MEM. If two MEMs are not overlapping or intersecting with each other, then a directed edge is constructed between them. For each node, the weight of each incoming edge is equal to the node weight. The CNSs are extracted from the graph using the approach similar to the longest path algorithm (Ma and Deogun, 2010). Here we use maximum flow algorithm instead of longest path for the weighted directed acyclic graph to find the path with maximum weight. The algorithm finds the optimal path consisting of CNSs that gives maximum cumulative score (Shown in Figure 1B). The ties are broken based on the distance between consecutive CNSs. If two or more CNS sets have same maximum score, then the set with minimum intra-CNS distance is chosen. In the Figure 1C, the first set of CNS is chosen among the four CNS set having equal score. The CNS that is selected among four have *min*{|*d*_1_ − *d*_2_| + |*d*_1_ − *d*_3_| + |*d*_2_ − *d*_3_|} where *d*_1_, *d*_2_ and *d*_3_ are the distances between two different MEMs in the sequence 1, 2 and 3 respectively. Basically, the CNSs with less variation in the distance between two of its consecutive MEMs is ranked higher.

The output of STAG-CNS is a file containing the list of CNSs with their start and end positions, length and the name of the chromosome. It also produces the visualization file that can be used in Gobe visualization tool (Pedersen et al., 2011). The source code for STAG-CNS is freely available at https://github.com/srbehera11/stag-cns.

### Genomes and Orthologous syntenic gene sets

The genomes and annotations of seven related species were downloaded from Phytozome (https://phytozome.jgi.doe.gov/pz/portal.html) (sorghum (Sorghum bicolor v3.1) McCormick et al. (2017), rice (Oryza sativa v7) Ouyang et al. (2007), setaria (Setaria italica v2.2) Bennetzen et al. (2012), Brachypodium (Brachypodium distachyon v3.1) Vogel et al. (2010), maize (Zea mays) Schnable et al. (2009)) and CoGe (https://genomevolution.org/coge/) (Oropetium (Oropetium thomaeum v1.0) VanBuren et al. (2015), Dichanthelium (Dichanthelium oligosanthes v1.001) Studer et al. (2016)). A pan-grass orthologous gene list including all these 7 species generated on the GEvo panel of CoGe is available (https://genomevolution.org/coge/Gevo.pl) (Schnable et al., 2016). A set of 200 orthologous syntenic genes of 6 species (excluding maize) were randomly selected from the pan-grass syntenic gene list and these were used to test the optimal minimum length of CNS when comparing orthologous genes from different numbers of species (Table S1). Another set consisted of 17,996 orthologous syntenic genes present in three species (sorghum, setaria, and rice) were also extracted from the pan-grass syntenic gene list (Table S2). A sub-set of 200 syntenic genes, which had the greatest number of CNSs between sorghum and rice based on CDP, was extracted (Table S3). These syntenic gene sets were used for the comparison of STAG-CNS and CDP. A set of 6,156 maize duplicated genes from two maize sub-genome and corresponding syntenic genes in sorghum and setaria were obtained from the pan-grass syntenic genes list (Table S7).

### Extracting non-coding regions associated with genes and permutation testing of CNS false discovery rates

The region used for CNS discovery started 10 kb upstream and ended 10 kb downstream pf a given syntenic gene. However, the sequence was truncated at the next conserved syntenic gene if this gene was less than 10 kb away from the the start or end of the target gene. For permutation testing, these sequences were grouped randomly across species for each permutation. The average number of CNS identified across 100 permutations was compared to the original set of CNS identified without shuffling syntenic relationships to estimate the false discovery rate. The difference between the average number of CNS from permutation test and the number of CNS from the original set (without shuffling) is considered to be false positives. The estimated true positive number of CNS was the number of CNS in syntenic genes minus the false discovery number of CNS. The true positive rate is ratio of true positive number of CNS and all number of CNS in syntenic genes. This analysis was repeated separately for minimum CNS lengths between 8 bp and 22 bp and for genes extracted from between 2 species and 6 species. The species compositions of each group were: two species (sorghum and rice), three species (sorghum, rice, and setaria), four species (sorghum, rice, setaria, and Oropetium), five species (sorghum, rice, setaria, oropetium, and brachypodium), and six species (sorghum, rice, setaria, oropetium, brachypodium, and dichanthelium).

### Comparing STAG-CNS and CDP using sorghum, rice, and setaria

The approach described in the previous subsection was also applied to extract the non-coding sequences 10 Kb upstream and downstream of the genes. A total of 17,996 syntenic genes of sorghum, setaria, and rice were used to find the CNS with optimal minimum length of CNS at 12 base pairs.

At the same time, the CNS in syntenic genes of sorghum-rice and sorghum-setaria were identified using CDP with default parameters (Turco et al., 2013) but with the modification that syntenic gene pairs were provided externally from the same list used by STAG-CNS rather than identifying syntenic gene pairs automatically (the default of the CDP). Then the CNSs of sorghum-rice and sorghum-setaria were combined based on positions on the genome of sorghum. The overlapped region (⩾ 12bp) of CNSs between two pairwise species were regarded as the CNSs of pan-grass syntenic genes. Summary statistics were computed with custom perl scripts and basic functions in R software.

### GO terms enrichment analyses

The enrichment analyses were performed using the goatools (version 0.5.9) (Haibao et al., 2015), a python script to find enrichment of GO terms. The genes in GO annotation file of sorghum was used as association data. A total of 17,996 sorghum syntenic genes were used as a reference population. Non-annotated genes were temporary annotated as a GO term that was not present in the analyses to test if these unannotated genes were enriched in any groups. The genes with 0,1,2, and ⩾; 3 CNSs were grouped respectively into four subsets. The Bonferroni correction and false discovery rate (fdr) implementation using re-sampling method were applied with significance p-value of < 0.05. The GO enrichment data for sorghum are shown in (Table S6).

### Association analyses between CNSs and DNase I sites

The DNase I sites of seeding and callus were downloaded from the NCBI GEO (http://www.ncbi.nlm.nih.gov/geo/) which have been generated by Zhang et al. (Zhang et al., 2012). The overlapping positions between DNase I sites and CNSs identified from 17,996 syntenic genes in sorghum, rice (v5), and setaria were calculated using custom perl scripts. A total number of 1,873 syntenic genes in 6 species (sorghum, rice, setaria, oropetium, brachypodium, and dichanthelium) were extracted from the syntenic gene list. The CNSs with minimum length from 8 bp to 22 bp are extracted from different numbers (2 to 6) of species identify the CNSs and compared with the DNase sites. The percentages of overlap of CNS and DNase sites for each case were calculated and shown in the heatmap (Figure 3C-D).

### Comparison of CNSs in maize sub-genomes

The set of 6,156 syntenic genes form sorghum, setaria, and duplicated maize genes (Table S7) were divided into two groups (sorghum-setaria-maize I and sorghum-setaria-maize II) that were used to identify the CNSs in the non-coding sequences 10 Kb upstream and downstream of these syntenic genes. The numbers of CNS were counted for each syntenic gene of the two groups and the differences between the duplicated genes, maize I and maize II, are computed using in-house perl scripts. The expression values of these duplicated genes in multiply tissues were calculated from several dataset Zhang et al. (2017); Chang et al. (2012); Bolduc et al. (2012); Waters et al. (2011); Wang et al. (2009); Chettoor et al. (2014); Davidson et al. (2011); Li et al. (2010) to compared the expression level between maize I and maize II. For case 1, maize I genes having more CNS than maize II, we count the number if maize I genes have higher/lower expression than maize II and calculate the ratio between them to measure the bias. For case 2, maize I genes having less CNS than maize II, we also count the number if maize II genes have higher/lower expression than maize I and calculate the ratio.

## Author contributions

J.C.S. and J.S.D. designed the research. S.B. and J.S.D. conducted the software development. X.L., S.B., and Z.L. tested the software and analyzed the data. X.L. and S.B. wrote the initial draft. All authors reviewed and edited the paper.

## Acknowledgments

This work was supported by internal funding to J.C.S. and J.S.D., and by a China Scholarship Council fellowship awarded to X.L.

## Supplemental Information (SI)

**Figure S1.**
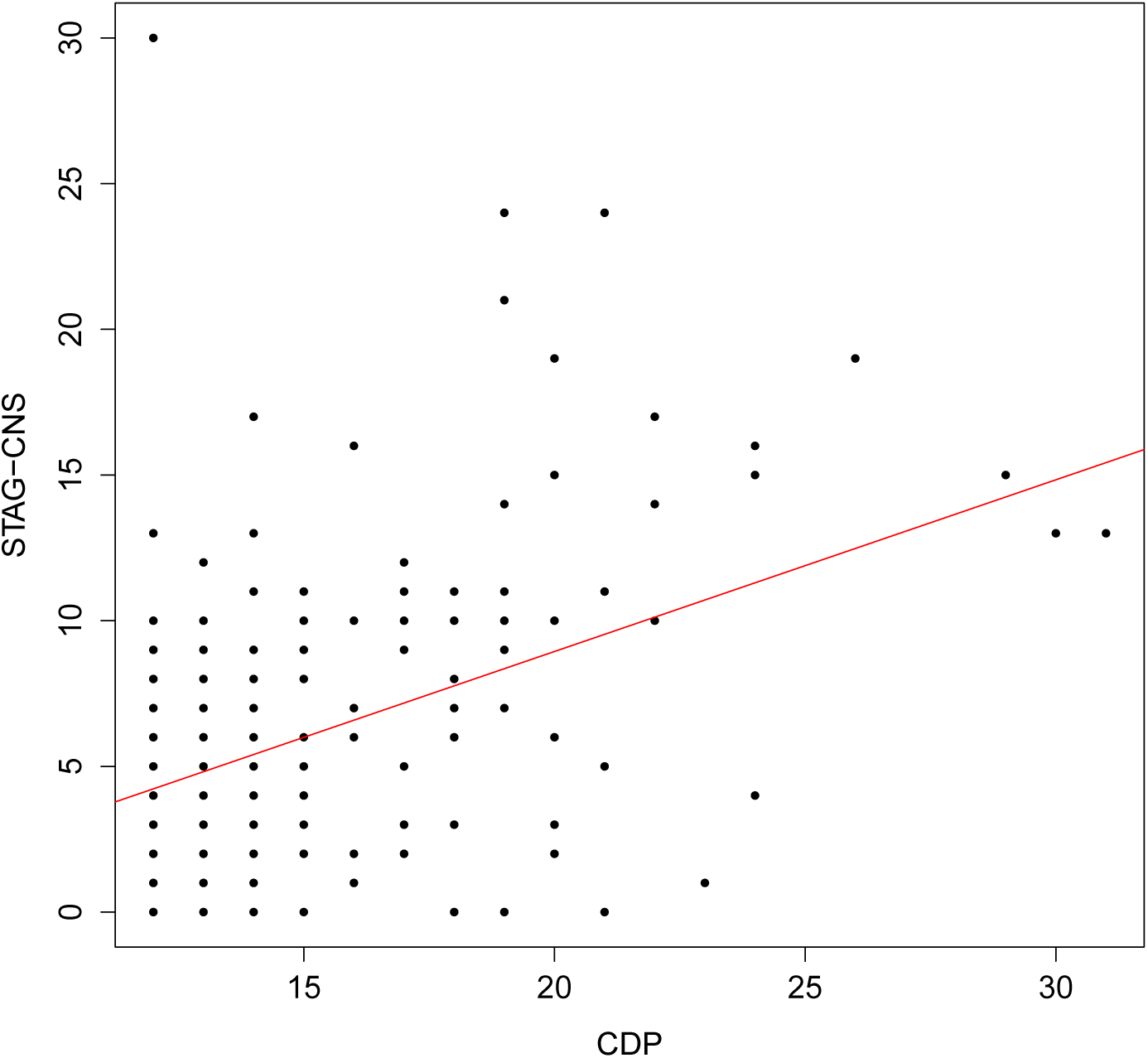
Correlation between the number of CNS identified by the CDP for a given gene and the number of CNS identified by STAG-CNS for the same gene. Red line marks the linear regression between the two datasets.

**Figure S2.**
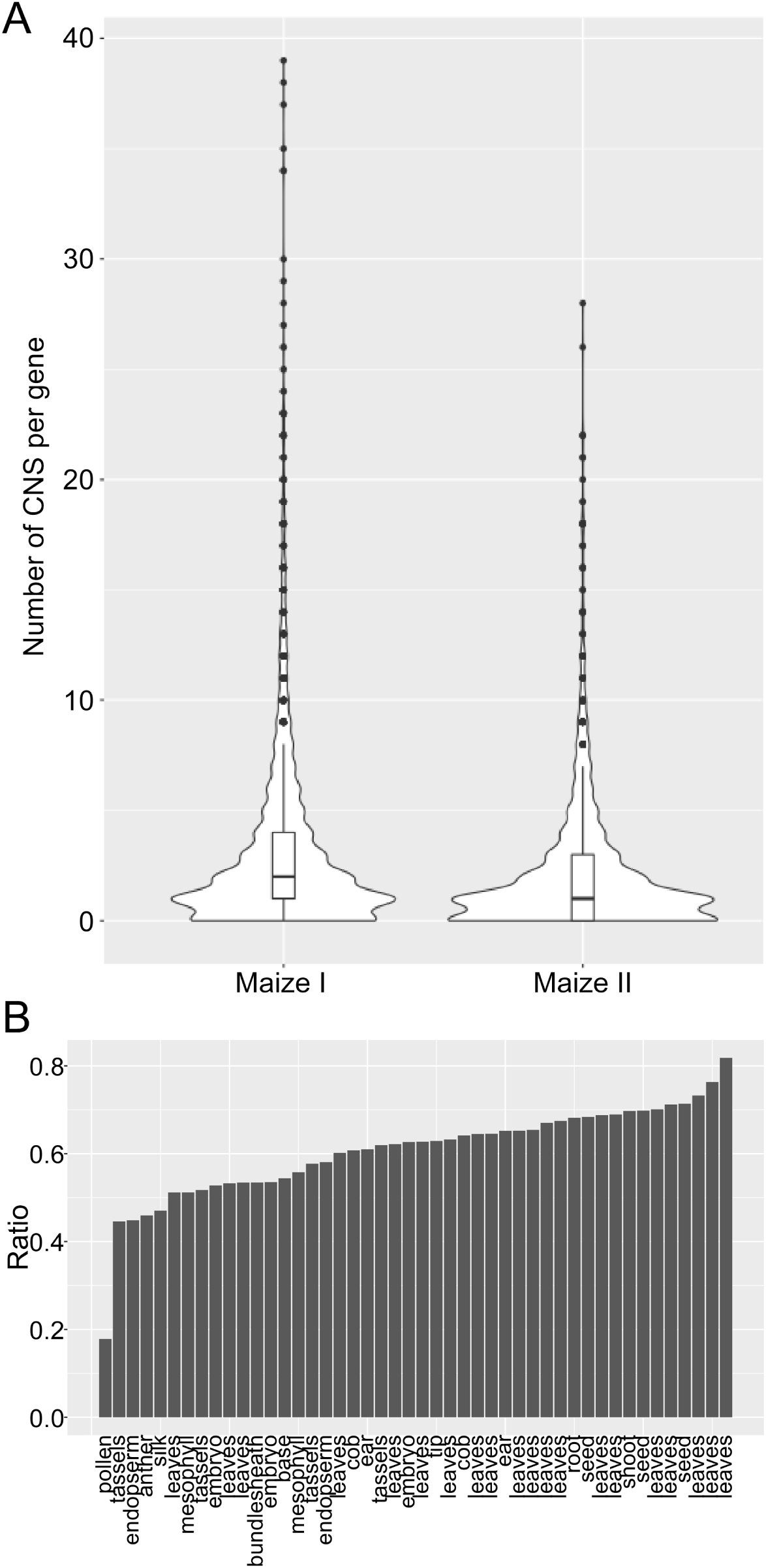
Statistics of CNS number and genes expression bias in two maize sub-genomes.(A) Boxplot showed the distribution of number of CNS in each retained gene. (B) Ratio of genes expression Bias between the cases that maize I retained more CNS and maize II retained more CNS in different tissues.

**Table S1**. List of the 200 random selected syntenic genes in 6 species used for initial permutation testing of the STAG-CNS algorithm.

**Table S2**. 17,996 orthologous syntenic genes in sorghum, setaria, and rice used for comparison of STAG-CNS to the CDP.

**Table S3**. The 200 rice sorghum syntenic gene pairs with the greatest number of associated CNS identified by CDP.

**Table S4**. The CNS identified by STAG-CNS using syntenic orthologous genes in sorghum, setaria, and rice.

**Table S5**. The distribution of CNS identified among sorghum, rice, and setaria on rice genome (reference genome v5).

**Table S6**. Results of GO enrichment analyses of sorghum genes associated with different numbers of CNS.

**Table S7**. Number of CNSs identified by STAG-CNS using syntenic genes conserved across sorghum, setaria, and maize I as well as equivalent results for syntenic genes conserved across sorghum, setaria, and maize II

